# Intrinsic fitness differences outweigh environmental matching in shaping introduction outcomes in nature

**DOI:** 10.64898/2026.02.04.699496

**Authors:** Lucas Eckert, Daniel I. Bolnick, Alison M. Derry, Grant E. Haines, Alexis M. Heckley, Åsa J. Lind, Catherine L. Peichel, Allison M. Roth, Natalie C. Steinel, Kristen Vlahiotis, Jesse N. Weber, Andrew P. Hendry, Rowan D.H. Barrett

## Abstract

Species introductions and transplants offer powerful contexts to understand evolutionary patterns and processes, and they are increasingly critical for conservation. However, introduction success varies widely, and predicting outcomes remains challenging. Introducing multiple source populations should increase the chance of success, while also providing an opportunity to explore the factors that predict success of individual source populations in the same environment. We used replicated, mixed-population introductions of >12,000 threespine stickleback (*Gasterosteus aculeatus*) to test whether source population success could be predicted by environmental matching between source and recipient environments and/or by intrinsic source population characteristics. We introduced four to eight source populations of stickleback into each of nine natural lakes and tracked their relative success over the following two years (up to two generations). Source population success was largely consistent across lakes, despite divergent environmental conditions. These results point to the importance of intrinsic source population characteristics rather than environmental matching in predicting introduction success in natural settings. Source populations that were consistently successful tended to have greater stress tolerance (mortality rate during translocation) and higher genetic diversity, though these relationships were not conclusive. Our study highlights the value of considering factors that generate fitness differences independent of environmental contexts in predicting ecological and evolutionary dynamics and planning conservation programs.

**Significance Statement:** Predicting which populations will succeed when introduced to new environments is a central challenge in ecology, evolution, and conservation. Environmental conditions are often assumed to be crucial in determining which populations succeed, given the expectation that populations preadapted to local conditions should perform best. We test this assumption by introducing >12,000 threespine stickleback fish from up to eight source populations into nine natural lakes spanning diverse environmental conditions. We show that the relative success of individual source populations was remarkably consistent across lakes and environmental conditions, indicating that some populations are intrinsically better suited to introductions. Our findings underscore the importance of considering intrinsic population characteristics alongside environmental conditions when predicting ecological and evolutionary outcomes and guiding conservation efforts.

## Introduction

Transplants and introductions have long served as a tool to understand evolutionary patterns and processes (1). Most notably, reciprocal transplants are often used to test the prevalence and strength of local adaptation, with varying conclusions (2, 3). Surprisingly, in numerous cases, non-local individuals appear to have higher fitness (i.e., survival/reproduction) than local counterparts, and a number of mechanisms have been proposed to explain such cases, such as gene flow constraining adaptation or limited adaptive variation (4, 5). Yet it remains unclear in which contexts such mechanisms can hinder or obscure local adaptation. The challenge of predicting fitness in particular environments becomes more complex when trying to extrapolate patterns of local adaptation beyond the current locations (or environmental niche spaces) occupied by the populations in question (6, 7). Specifically, do populations perform best in environments that resemble their native habitat? Poor performance of these extrapolations limits our ability to predict population persistence or success in several critical scenarios; for example, the establishment/spread of invasive species (8), and forecasting population declines (7, 9) or range shifts (10) under climate change.

Similar predictive challenges arise in conservation, where introductions or translocations (e.g., restoration or assisted colonization/migration) are becoming an increasingly important tool (11–13). In many cases, however, such initiatives fail to establish new populations or achieve only marginal success (14). Indeed, the success rates of introduction programs have not improved over time (15), and the factors influencing establishment and growth of specific introduced populations remain poorly understood (16–18). Still, two consistent recommendations have emerged: first, to maximize the number of introduced individuals (16, 19, 20), and second, to maximize their genetic diversity (21, 22). Increasing the number of individuals intensifies propagule pressure (23, 24), while greater genetic diversity reduces the chance of inbreeding depression (25), increases the likelihood that some genotypes are initially well-adapted, and enhances long-term adaptive potential (26–28).

Mixing multiple source populations efficiently increases genetic diversity (29), and there is growing support for this approach in conservation efforts (30, 31). Introduction programs that release multiple source populations together into a given location also provide opportunities to test which factors drive success of individual source populations. Two general hypotheses have been advanced. First (H1), assuming local adaptation is both prevalent and strong, populations from environments similar to the recipient site should be more successful (32, 33). Indeed, this principle underlies the common recommendation of environmental matching for conservation programs (31, 34, 35). Second (H2), some populations might possess intrinsic advantages independent of environmental context. For example, greater genetic diversity (36, 37) or faster growth rates (38) for particular populations might confer advantages irrespective of the environment into which they are introduced. These intrinsic advantages could override environmental effects, making some source populations consistently more successful. Importantly, these hypotheses are not mutually exclusive; both could be at play and the relative importance of each may shift with changes in demography, genotype frequencies, or environmental conditions. In the present study, we explore how the above two hypotheses predict source population success in natural environments through replicated introductions of multiple threespine stickleback (*Gasterosteus aculeatus*) populations.

Threespine stickleback (hereafter stickleback) are an important system in evolutionary biology, largely due to their propensity to colonize and adapt to new environments (39), often resulting in striking patterns of phenotypic divergence (40). These patterns of divergence often reflect local adaptation, and are seen in strong and repeatable phenotype-environment correlations (41–43). One such pattern is seen in variation among lake stickleback, where individuals and populations can be categorized along a continuum of *benthic* to *limnetic* ecotypes (44–47). Divergence of these ecotypes has been primarily linked to lake bathymetry, likely through its influence on prey community structure and abundance, which in turn influences the trajectory of stickleback adaptation (46, 47). As a result, *benthic* fish are typically found in small shallow lakes, where they feed predominantly on benthic macroinvertebrates, whereas *limnetic* fish are typically found in large deep lakes where they feed on limnetic zooplankton – with well-defined phenotypic differences between ecotypes (44, 46). Although particularly well-established in stickleback, similar patterns of strong local adaptation are found in other commonly introduced species, including salmonids and sunfish (48, 49).

Given that environmental conditions drive striking patterns of adaptive divergence in stickleback, environmental matching (H1 above) could be critical in determining the relative success of introduced source populations. At the same time, certain population-level characteristics (H2), such as genetic diversity, body size, stress tolerance, and parasite load also vary dramatically among stickleback populations, including within ecotypes (50–52). These population-level differences could account for fitness differences independent of environmental conditions, making some populations consistently more successful. Here, we evaluate the relative support for these hypotheses by examining multi-generational success of multiple populations sourced from diverse environments in replicated, whole-lake introductions. Much of the previous work on this topic has relied on experiments in controlled settings (e.g., competition trials in laboratories or mesocosms) or post-hoc analyses of conservation introductions or biological invasions, which are respectively limited in their extension to natural settings and their ability to capture population dynamics as they unfold. As such, this study provides valuable insight into the dynamics of population success in real-time and in complex natural environments.

## Results

### Experimental design and implementation

To test the predictors of source population success, we used replicated, mixed-population stickleback introductions into natural lakes for the purposes of restoration. In 2019, we introduced stickleback to nine fishless lakes in the Kenai Peninsula of Alaska (Figure 1), hereafter referred to as the recipient lakes. These lakes were treated with rotenone the previous year by the Alaska Department of Fish and Game to eliminate invasive predatory fish, which also eliminated all other fish species (53). We stocked each of these lakes with stickleback drawn from either four nearby benthic source lakes, four nearby limnetic source lakes, or a mixture of both ecotypes from all available source lakes, scaling the number of individuals non-linearly by lake area, while achieving a relatively equal mixture of fish from each source (Figure 1; Table S1 for lake details; Table S2 for stocking numbers). While the source populations were selected based on the morphology of the fish, the pairings of source ecotype to recipient lake were chosen to ensure that each ecotype treatment was introduced into lakes of varying environmental and ecological conditions, as illustrated by relative littoral area in Figure 1. See Hendry et al. (54) for further details on the design and implementation of the experiment.

**Figure 1:**
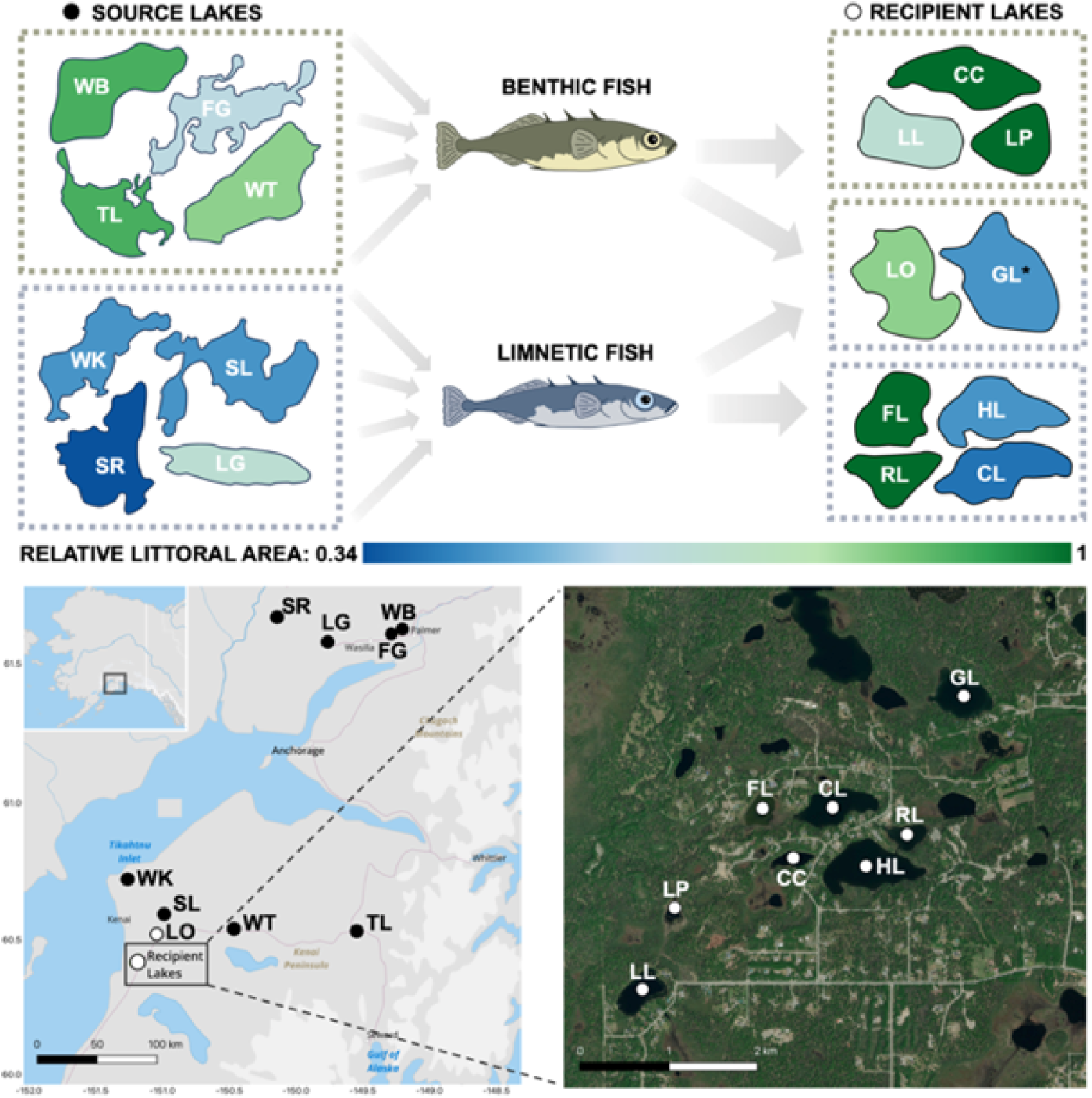
Experimental design: Lakes are coloured according to relative littoral area (with green being shallower than blue) to illustrate the range of environmental similarity among source and recipient lake pairs. The maps show locations of source (black) and recipient (white) lakes with reference to nearby landmarks.

All but one of the original introductions were successful. We originally stocked G Lake (GL) with only benthic fish, but it quickly became apparent that the introduction failed, as no fish were caught in extensive sampling efforts over the two years following the introduction. In 2022, we restocked with both benthic and limnetic fish, although one of the source populations (Long Lake: LG) was not used due to low abundances, so GL only received seven of the eight source populations – and the second attempt succeeded. The original introductions succeeded in the other lakes, where populations appeared to grow rapidly; anecdotally, catch rates were comparable or higher than those in the source lakes just one year post-introduction.

### Predictors of source population success

To measure the differential success of each introduced source population, we sampled 100 fish from each recipient lake one year and two years post-introduction and genotyped those fish to infer source population ancestry. Ancestry was assigned using 158 single nucleotide polymorphisms (SNPs), each of which contained an allele unique to a single source population and found in high frequency within that population (20 SNPs unique to each source population on average, with a mean frequency of 0.81; details in Materials and Methods). Most of the fish were admixed in these post-introduction generations (72% in the first year and 80% in the second). As such, we estimated the proportional ancestry of each source population in each fish, rounding to the multiples of 0.5 in year 1 and 0.25 in year 2, as the genotyping data suggested a minimum generation time of just one year. To quantify the “success” of each source population in each lake in a given year, we summed the proportional ancestries across the sample of fish, giving us the overall proportion of ancestry from each source population in each lake. We then analyzed how those proportions changed over time in each lake and whether those changes were predicted by environmental similarity between the source and recipient lakes or intrinsic characteristics of the source populations.

Large shifts in proportions of source population contributions were seen in each lake in the first two years post-introduction (Figure 2). These shifts often exceeded those expected by drift and sampling error – as assessed with simulations (Figure S4; Table S4). We analyzed variation in these changes in proportions across three different timespans: *year one change*: the change between the initial stocking proportions and proportions one-year post-introduction, *year two change*: the change between one- and two-years post-introduction, and *net change*: the change between initial stocking proportions and proportions two years post-introduction. Using linear mixed-effect models, we tested if the observed changes in proportions over each of these timespans (analyzed as log-ratios) were predicted by environmental distance between the source and recipient lakes (H1), and/or source population identity (H2). We also included the initial proportion of each source population over that timespan as a fixed effect to account for the possibility of frequency-dependence, though this term was insignificant in all models (Table S5).

**Figure 2:**
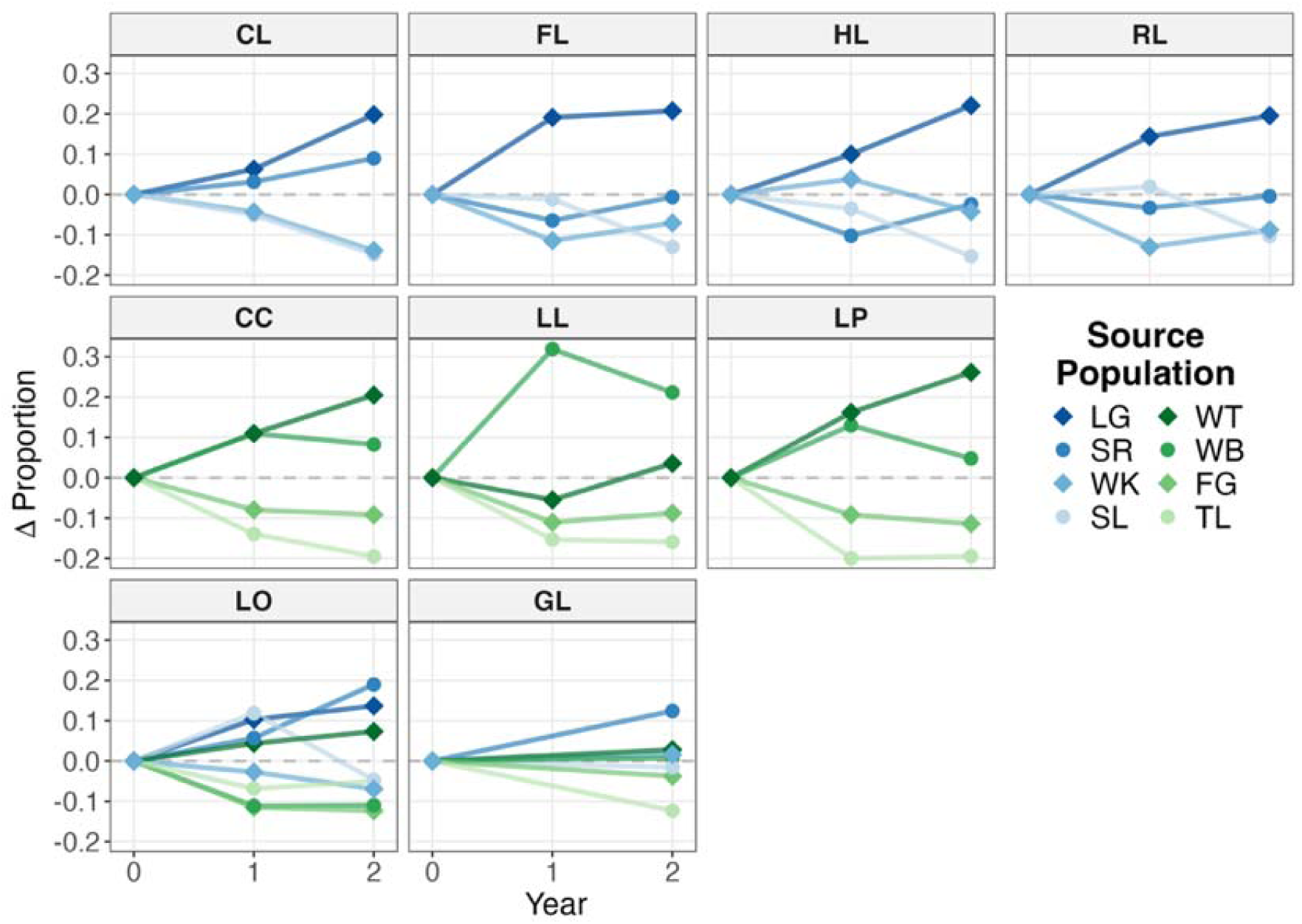
Source population success: The change in proportion of each source population in each recipient lake with respect to initial stocking proportions. Limnetic source populations are shades of blue and benthic are green (different shapes are used for clarity). Trends in proportions not adjusted to initial stocking proportions are shown in Figure S5.

We computed individual models for each timespan and for three different metrics of environmental distance (nine models total). Briefly, the metrics of environmental distance ranged from a simple set of variables (metric 1: relative littoral area and calcium concentration, both of which have well-documented associations with stickleback adaptive divergence) to an intermediate set (metric 2: adding abundances of important prey taxa) to a complete set (metric 3: all measured environmental variables plus prey abundances). We found that environmental distance (H1) did not explain the observed changes in source population proportions, and that this conclusion held for all three timespans and across all three metrics of environmental distance (Table S5). In contrast, changes in proportions were well predicted by source population identity (H2), as that term was significant in all but one model, where it was marginally significant (Table S6) – indicating that some source populations were consistently more successful than others (Table S7).

### Temporal dynamics of population success

When investigating the mechanisms underlying changes in source population success, additional insight can be gained by examining the timing of those changes. The outcomes of these introductions might be strongly influenced by processes operating over very short timescales, such as days or weeks after introduction, or by longer-term dynamics that persist beyond the timeframe of our study. In the latter case, substantial fluctuations beyond the first two years could indicate the patterns observed in the timeframe of our study might not be indicative of the final outcome of the system. To explore the chronology of changes in source population proportions, we conducted additional sampling on shorter and longer timespans than those represented in our primary analysis. For one of the lakes that received both ecotypes (GL), we sampled one month after introduction and observed significant shifts in source population proportions (χ^2^ = 18.41, df = 6, p = 0.005) similar in magnitude to the changes observed after two years (Figure 3A), indicating that shifts in population proportions can occur quite rapidly. For the other lake that received both ecotypes (Loon Lake: LO), we sampled the population for an additional year (three years post-introduction) to explore if these lakes were still experiencing large shifts in source population proportions. The dynamics observed in this lake indicate that the magnitude of change generally decreased as time went on (Figure 3B; F = 8.025, df = 22, p = 0.01). By the third year, the changes in source population proportions were ∼4X smaller in magnitude on average than those in the first year.

**Figure 3:**
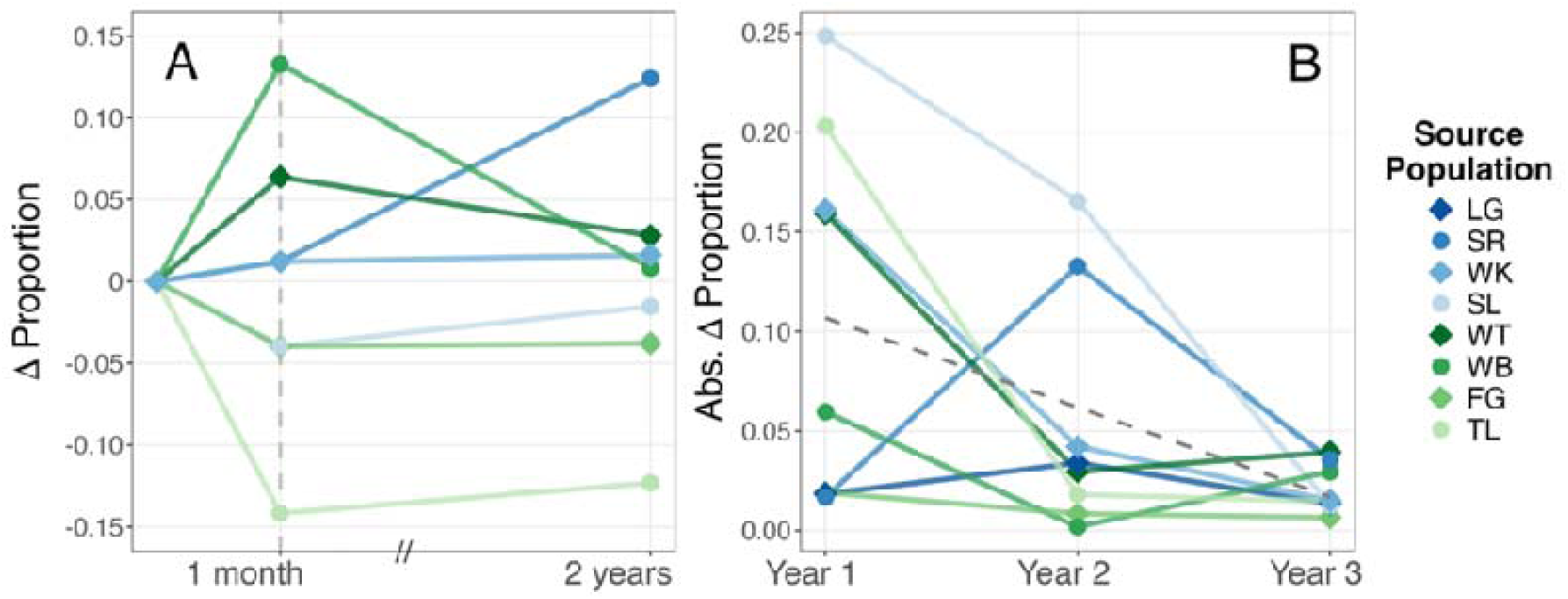
Short- and long-term trends: **(A)** Changes in proportions of each source population after one month in G Lake (GL), compared to the subsequent changes observed after two years. **(B)** Absolute magnitude of change in proportions from the previous year for each source population in Loon Lake (LO) across the first three years.

### Characteristics of successful populations

As some populations were consistently more successful (H2), we tested to what extent population-level characteristics could explain this pattern. To do so, we regressed the mean estimated marginal effect size of each source population (averaged across our three models of different measures of environmental distance), against population means of the following population characteristics: median body size, mortality rate during translocation (stress tolerance), parasite (*Schistocephalus solidus*) prevalence, and genetic diversity (Figure 4; Table S8). Statistical power was limited (only eight populations) and, as such, this analysis was intended more to be exploratory than definitive – the main goal being indication of variables that could be investigated in future studies. Mortality rate during translocation (explained in Materials and Methods) had the strongest effect on year one success (β = -0.665), which was marginally significant before correcting for multiple testing (p = 0.072; p_adj_ = 0.288). Whether or not mortality rates during translocation influenced subsequent population success, mortality rates varied significantly across source populations (Figure S3), which has practical implications for source population selection in conservation contexts. The predictor with the next strongest effect was genetic diversity in year one (β = 0.487), although this effect was not significant (p = 0.221; p_adj_ = 0.442).

**Figure 4:**
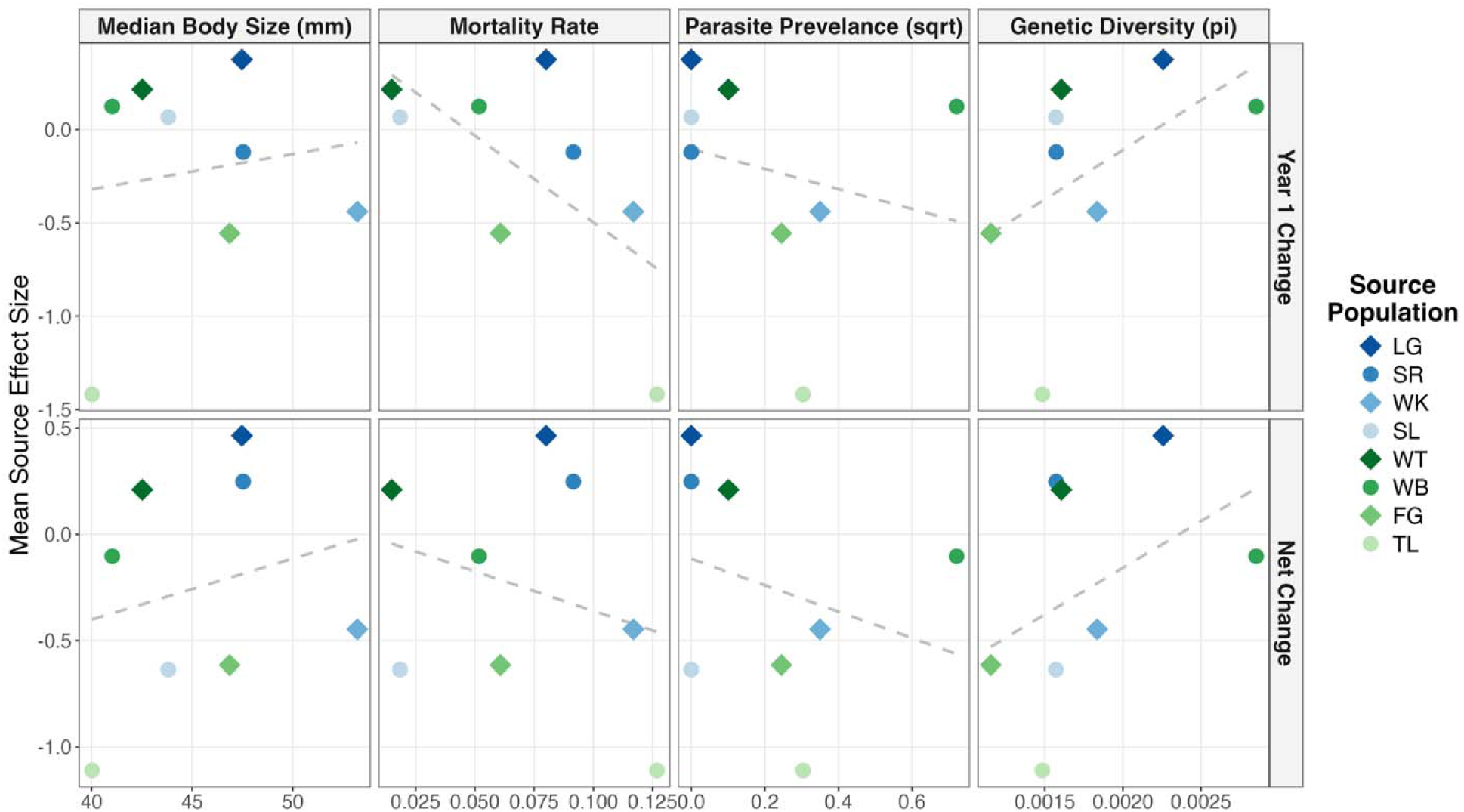
Characteristics of successful populations: Estimated marginal mean effect size of each source population (averaged across the three measures of environmental distance), for both year one change and net change in source proportions, correlated against several population-level characteristics. Dashed lines are individual linear models, none of which were statistically significant after correcting for multiple testing (Table S8).

## Discussion

We documented substantial variation in source population success following replicated introductions into new environments. These differences in success arose quickly and resulted in dramatic differences in source population proportions after just two years. Overall, the data strongly support the hypothesis that intrinsic differences between source populations predict their relative success and provide no support for the hypothesis that environmental matching between source and destination habitats plays a crucial role in success.

The lack of environmental matching in our study is in contrast to considerable evidence from prior studies in which environmental matching predicts success in similar scenarios (34, 55). For example, climate suitability has been linked to the success of conservation translocations in plants (56, 57), and evidence from invasion ecology suggests that environmental matching often contributes to the spread and impact of invasive species (58, 59). The contrasting lack of support for environmental matching in our study could be explained by several factors. First, many of the studies that investigate environmental matching measure persistence of single source populations rather than competitive outcomes among multiple populations. The environmental conditions in all of the recipient lakes are likely well within the range required for the persistence of all source populations – and, indeed, historical records indicate that stickleback were previously present in all of these lakes before their elimination by invasive predatory fish (60). Moreover, stickleback are generally able to persist across a wide variety of environmental conditions, as evidenced by their extensive history of colonization and range expansion (39). Therefore, selection is likely acting on traits related to intraspecific competition and population growth – as opposed to abiotic stressors. Although competitive ability can still be influenced by the environment (e.g., trophic traits related to interactions with prey community), these differences might be swamped by fitness differences between populations that are largely unrelated to the environment (e.g., life history traits). This highlights the importance of considering factors that contribute to fitness that are both dependent and independent of the environmental context (61). Indeed, conceptual challenges in studies of local adaptation often arise from an emphasis on genotype-environment interactions while overlooking intrinsic fitness differences among genotypes (62).

The potential importance of competition in shaping our results suggests that environmental matching may become more influential over longer timescales, as selection from intraspecific competition intensifies with increasing population density. However, this would only occur if the resulting selection acts on traits that interact with the environment (e.g., trophic traits). Further, results from a third year of sampling in Loon Lake (LO) show that the magnitude of changes in source proportions are decreasing over time (Figure 3B). In essence, source population frequencies shifted quickly at first but might begin to stabilize in later generations. This can be partially explained by the high level of admixture observed in our populations (80% of fish were admixed in year 2); as the level of admixture increases, fitness differences will be better predicted by individual genotypes rather than source population ancestry (63). Finally, early ecological and evolutionary dynamics often have a disproportionate influence on long-term trajectories and outcomes – as is the case in community assembly (64, 65) and adaptation (66–68) – suggesting that environmental matching might only be expected to have a strong influence on long-term outcomes if it was manifested in the generations captured by this study.

Although some populations were consistently more successful than others, it is not clear what mechanisms drove those differences. Stress tolerance (translocation mortality in our case) and genetic diversity had the largest effect sizes in our models, suggesting they might contribute to variation in source success, though these effects were not significant. Still, in support of these associations, evidence from invasion ecology shows that stress and thermal tolerance are commonly associated with invasive ability (69–71). Similarly, it has been noted that considering stress tolerance is critical in maximizing the success of conservation translocations (72–74), as these are inherently stressful events, resulting in both immediate mortality as well as lasting reductions in survival and fitness post release (75, 76). Greater genetic diversity is another characteristic theorized to be advantageous in these scenarios. Genetic diversity has been linked to establishment success in invasion ecology (77, 78), and maximizing genetic diversity is a common recommendation in conservation efforts (22, 27), as it is generally thought to be associated with fitness (79). Aside from stress tolerance and genetic diversity, other likely candidates to explain variation in success would be traits that contribute directly to population growth, such as growth rate or fecundity (80, 81). Although our proxy for these traits (median body size) did not predict variation in success, it is possible that growth rate and fecundity would if measured directly. Finally, it is possible that some advantages were conferred by plasticity or persistent carryover effects from the source environment rather than intrinsic characteristics.

Regarding conservation efforts, the sheer variation in source population success in our study indicates that the choice of source populations could be crucial in the outcomes of introduction programs. By contrast, environmental matching between the source and recipient habitats might not be a good predictor of population fitness. The absence of any effect of environmental matching is particularly noteworthy in our case, as we had detailed environmental and ecological data on variables with known associations with stickleback adaptation, including in our own source populations (47, 54, 82) – most introduction programs are unlikely to have such knowledge. Yet even in our case, extending those correlations to new habitats proved inaccurate. In choosing source populations, traits like stress tolerance and genetic diversity might be indicative of higher probabilities of success, but the importance of these traits can be context dependent (77). Therefore, we suggest that the most reliable way that conservationists can maximize the chance of success in introduction programs is to increase the number of individuals released and mix multiple source populations (30, 31). Using multiple source populations maximizes adaptive potential and, perhaps most critically, mitigates the risk of selecting source populations inherently less likely to succeed in these contexts, regardless of whether this is due to environmental matching or intrinsic characteristics.

An additional anecdote from our study also points to the utility of maximizing the number of source populations and individuals. Our first introduction attempt in G Lake (GL) in 2019 involved 1600 individuals from four benthic source populations. This attempt was unsuccessful as no fish were captured in subsequent years despite high effort. Our second introduction attempt into the same lake in 2022 used seven source populations (four benthic and three limnetic) and more than double the number of individuals (almost 3500). This second attempt was immediately successful, with high capture rates even in the first post-introduction year. Interestingly, the most successful population in this second GL introduction was limnetic (SR: South Rolly), and the environment of GL (low relative littoral area and calcium concentration) should suit limnetic ecotypes (47), suggesting that environmental matching could have been important in this specific context. However, the success could simply be due to the increased number of individuals and populations in the second attempt. Either way, the success of the second introduction exemplifies how introducing multiple source populations mitigates the chances of choosing populations that are ill suited to the destination environment or translocation contexts generally.

Local adaptation is far from ubiquitous and, in most cases, the precise connections between the environment, phenotype, and fitness remain poorly understood. As such, relying heavily on projecting trends of local adaptation across environmental space to predict fitness is a risky endeavor (6). Nevertheless, predicting fitness in new and changing environments is imperative to understanding and protecting biodiversity, with clear applications in predicting how species may respond to climate change (7), predicting the establishment and spread of invasive species (8), and planning conservation introduction programs (54). Our study highlights the importance of considering characteristics that generate fitness differences independent of environmental contexts. Such characteristics may continue to influence fitness even when environmental effects are present, and considering both mechanisms together could improve predictions of ecological and evolutionary dynamics and aid biodiversity management and conservation.

## Materials and Methods

### Experimental design

To explore the predictors of source population success, we use a replicated introduction experiment in natural lakes, the design and implementation of which are detailed in Hendry et al. (54) and summarized here. In 2018, nine natural lakes in the Kenai Peninsula of Alaska were treated with rotenone by the Alaska Department of Fish and Game (ADFG), eliminating invasive northern pike (*Esox lucius*) in eight lakes and invasive muskellunge (*Esox masquinongy*) in one lake, while also incidentally eliminating all other fish life (53, 60). In 2019, we reintroduced threespine stickleback (which are native and abundant in the region) to these lakes as part of an effort to rehabilitate these ecosystems, as natural colonization was prevented by bogs. Each of these lakes was stocked with stickleback drawn from either four nearby benthic ecotype source lakes, four nearby limnetic ecotype source lakes, or a mixture of both ecotypes from all available source lakes, while attempting to achieve an equal mixture of fish from each source (Figure 1; Table S1 for lake details; Table S2 for stocking numbers). The pairings of source ecotype to recipient lake were chosen to ensure that each ecotype treatment was introduced into lakes of varying environmental and ecological conditions, though it should be noted that the source populations themselves were selected based on the morphology of the fish (i.e., where they fell on the benthic-limnetic continuum), not the environmental conditions of the lake (54). The number of fish introduced to each lake was scaled non-linearly by lake size, from 386-3495 fish.

As discussed, one of the two lakes that received both ecotypes is exceptional for the following reasons, again detailed in Hendry et al. (54). G Lake (GL) was originally stocked with just benthic fish, but after it became apparent the introduction failed, it was restocked with both benthic and limnetic fish in 2022; although one of the source populations (Long Lake: LG) was not used due to low abundances, so GL only received seven of the eight source populations. GL was also stocked with a greater number of individuals than the other lakes (roughly double the original stocking plan), to increase the chance of success.

### Environmental distance

We measured potentially relevant environmental and ecological variables in both the source and recipient lakes, including physical features (relative littoral area), abiotic measures (pH, calcium concentration, dissolved organic carbon, dissolved oxygen, temperature, Secchi depth, total nitrogen, total phosphorus, chlorophyll *a* concentration (a measure of phytoplankton biomass), and specific conductivity) and biotic measures (abundances of crustacean zooplankton and benthic macroinvertebrates taxa). Methods for measuring each variable are described in the Supporting Information and environmental divergence among lakes across all variables is shown in Figure S1. We used these data to compute five metrics of environmental distance between lakes as scaled distance matrices, including five different combinations of these environmental and ecological measures. We computed distance matrices and performed all subsequent analyses in R (v4.5.1).

1. *Relative littoral area (RLA) and calcium concentration*: While many environmental variables impact stickleback fitness and evolution, RLA and calcium concentration have been strongly implicated in patterns of adaptive divergence. RLA quantifies the proportional area of the lake classified as the littoral zone, which we define as having a depth of less than three metres (calculated from bathymetric maps). RLA is a good predictor of stickleback morphology (43, 51, 83), including in our source lakes (Figure S2A), likely through direct effects on swimming-related traits and indirect effects on trophic traits by structuring the prey community. Similarly, calcium concentration has been linked to stickleback morphology through correlational studies (84–86), experimental studies (87), and again in our source lakes (Figure S2B), as calcium influences prey community structure (88, 89) as well as morphological investment, being critical for bony structures in stickleback such as armour and gill rakers (87). The associations between these variables and stickleback morphology make them likely candidates for predictors of stickleback fitness.
2. *RLA, calcium, and prey abundances*: To more directly capture divergence in prey community, we included abundances of choice prey for stickleback. Specifically, we included the combined abundance of two groups of benthic macroinvertebrates (family Chironomidae and family Gammaridae) and the combined abundance of two groups of pelagic zooplankton (order Calanoida and *Daphnia spp*.). All of these taxa are important prey items for stickleback (90–93) and the combined abundance measures should effectively capture the levels of benthic and limnetic prey resources, making it particularly relevant to the ecological divergence among our source populations. We measured these prey abundances in the recipient lakes after the rotenone treatment, but before the stickleback were introduced, averaged over two years of sampling (2018-2019).
3. *All abiotic variables and prey abundances*: For a comprehensive metric of environmental divergence among lakes, we included all the abiotic variables listed above in addition to RLA and calcium concentration. This metric complements the previous more targeted metrics by capturing additional environmental variation, at the risk of including certain variables that have limited impacts on stickleback fitness. We included the previously described combined abundances of prey types in this metric as well.

### Sampling and DNA extraction

We sampled 100 fish from each recipient lake one year and two years post-introduction, in late May of each year, using unbaited Gee’s (G40 – ¼” mesh) minnow traps, set around a single consistent location in each lake. All lethal sample collection was conducted with IACUC approval (University of Connecticut: A19-015, A22-006) and permits from the Alaska Department of Fish and Game. We euthanized the fish on site with buffered Tricaine methanesulfonate (MS-222), measured standard length (to 0.01 mm) and mass (to 0.01 g), and took a tissue sample from the caudal fin for DNA extraction, which used a standard phenol-chloroform protocol. Tissue samples were incubated at 55°C in digestion buffer with proteinase K. We isolated the DNA using an isoamyl-phenol-chloroform solution and precipitated in ethanol, and we quantified each extract using PicoGreen before genotyping.

### Genotyping and ancestry inference

To quantify the success of source populations, we inferred the ancestry of fish sampled from the recipient lakes and estimated the total genetic contribution of each source population to the introduced populations. To infer ancestry, individual fish were genotyped for single nucleotide polymorphisms (SNPs) unique to each source population, identified through pool-seq data from Weber et al. (52); specifically, 24 SNPs unique to each source population (192 SNPs in total). We selected SNPs to maximize the frequency of the unique alleles in the respective source populations, while filtering out SNPs with low reads (more than 1.5 standard deviations fewer than the mean read number) and minimizing effects of linkage by ensuring that every chromosome (excluding the sex chromosome) had at least one SNP. We designed two Fluidigm SNPtype assays, one with SNPs from the benthic source populations and one with SNPs from the limnetic source populations. To validate the efficacy of these assays in correctly inferring ancestry, we genotyped 16 individuals from each source population from samples collected in 2018, all of which were assigned to the correct population based on the genotyping results. This trial run also identified SNPs that were unsuccessful or that underperformed (i.e. were present in the trial fish in much lower frequency than expected from the pool-seq data); these were omitted from subsequent analyses, leaving 20 unique SNPs per population on average. The final list of SNPs, 158 in total with an average unique-allele frequency of 81%, can be found in Table S3.

Using these assays, we genotyped 95-97 fish from each of the recipient lakes from both 1 year and 2 years post-introduction. To infer the ancestry of fish from the genotyping data, we computed an ancestry “score” for each source population for each individual. This score was based on the number of unique alleles identified from each source in that individual, weighted to account for differences in the number of SNPs included for each source population and the average frequency of those alleles in those populations. Finally, the ancestry proportion estimates for each individual fish were rounded to the nearest fraction deemed possible according to the maximum potential generations, while ensuring the proportions always summed to 1. We assumed the fastest generation time to be one year (which was supported by further analysis of the genotyping data); therefore, for each individual, ancestry proportions (contributions from each source lake) were rounded to the nearest multiple of 0.5 for the sample 1-year post-introduction and 0.25 for 2-years post-introduction.

To confirm the accuracy of these inferences, we simulated analogous scenarios in R, parameterized with the number and frequency of the SNPs in our assays. In the simulated F1 generation, we had an error rate (inferring an incorrect rounded proportion of ancestry for at least one population in an individual) of 0.005% across 1000 simulations, and in the F2, we had an error rate of 6.5%. For the individuals from the F2 generation whose ancestry we mis-infer, we still identify the correct set of ancestral populations (but infer at least one wrong proportion) about half the time, leaving just 3.5% of F2 individuals where we miss a contribution from one of the ancestral populations.

### Mortality during translocation

There was some mortality during the translocation process, 6.8% of fish in total. These mortalities occurred after fish from multiple populations were pooled to be transported to each recipient lake, meaning that we could not immediately determine the source population of the fish that died. To account for these mortalities, we genotyped these fish (621 in total) to infer their ancestry and adjusted the initial stocking numbers of each source population accordingly.

### Source population success

To quantify the success of each source population in each recipient lake, we summed the proportional ancestries across every fish in that lake. This calculation was made for each year individually, giving us the total genetic contribution of each source population to each recipient population in each of the first two years post-introduction; these measures are subsequently referred to as the “proportion” of that population in a given recipient lake at a given time. Since each lake started with slightly different proportions in the mixture of source populations (Table S2), we analyzed source success as changes in proportion between timepoints. To test if different mechanisms were operating over different time spans, we computed three different measures of changes in proportions: *year one change*: the change between the initial stocking proportions and proportions one year post-introduction, *year two change*: the change between one and two years post-introduction, and *net change*: the change between initial stocking proportions and proportions two years post-introduction.

We first explored if the observed changes in proportions were beyond those expected under neutral conditions; specifically, drift during exponential population growth and sampling error. In R, we simulated drift and sampling error in each of our lakes, parameterized with the actualized stocking numbers of each source population and assuming a high population growth rate of 50 offspring per adult fish; this growth rate was chosen as it is likely towards the upper end of biologically realistic values, reducing the chance that we are underestimating the effect of rapid population growth. In 1000 replicates of each scenario, we simulated two generations of drift during population growth, followed by random sampling of 100 fish per lake, analogous to the sample of fish that we genotyped after year two. We calculated the change in proportions of each source population and compared those null distributions to the observed changes in proportions (Figure S4). Statistical significance was determined by the proportion of simulated values with an absolute magnitude greater than the observed change (Table S4).

We then used linear mixed-effect models (LMMs) to test predictors of source population success, change in proportions of each source population in each lake as the response variable. Changes in proportions were analyzed as log ratios (the natural logarithm of the ratio of final to initial proportions), to reduce boundedness and compositional constraint. We added a small pseudo-count (equal to the smallest detected proportion in our dataset) to both the initial and final proportions, to account for proportions smaller than the detection threshold (which we would incorrectly measure as zero). We included environmental distance (discussed above), source population identity, and initial proportion (for that time span) as fixed effects. Including environmental distance allowed us to test if environmental matching predicts population success, while source population identity allowed us to test if some source populations are consistently more successful. Finally, including the initial proportion of that source population in that time span allowed us to test for frequency dependence. To further account for the non-independence of proportion measures from the same recipient lake, we added recipient lake as a random effect. As we modelled measures of success over three different timespans with three alternative measures of environmental distance, this resulted in nine models (Table S5). We fit models through *lme4* (v1.1-37) and estimated standardized effect sizes and p-values of fixed effects with the *effectsize* (v1.0.1) and *lmerTest* (v3.1-3) packages. To account for multiple testing, all p-values were adjusted using a Benjamini-Hochberg correction. Type III F-tests were used to estimate the significance of the source identity term.

### Short-term and long-term trends in success

To better understand the chronology of changes in source population success, we conducted additional sampling on shorter and longer timespans. For one of the lakes that received both ecotypes (GL), we sampled the population just one month post-introduction. Although this only represents a single replicate, thus limiting our ability to generalize the findings, this analysis does provide insight into the timescale in which these differences in success arise. Roughly one month (31-37 days) post-introduction, 58 fish were sampled using minnow traps from two locations on the lake, one close to the point of release and one on the opposite side of the lake. Tissue samples from these fish were genotyped to infer ancestry and assign a source population (Figure 3A). To test if the observed number of fish from each source population significantly diverged from our null expectation of random survival, we used a chi-square goodness-of-fit test.

For the other lake that received both ecotypes (LO), we sampled the population for an additional year (3 years post-introduction). In this case, 100 minnow traps were set around the perimeter of the lake, and two fish were collected at random from each trap, resulting in 200 fish being genotyped. Again, while this addition is only a single replicate of its sampling type, these data provide insight as to whether the population was still experiencing large shifts in source population success or if the proportions of each source population seemed to be stabilizing (Figure 3B).

### Characteristics of successful populations

To test if trait differences predict which populations were most successful, we compared the mean success of each source population against the following characteristics that might explain differential survival or competitive ability: mortality rate during translocation, median body size, parasite prevalence, and genetic diversity. Mortality rates might be predictive as stress effects from translocation might have led to additional mortality or latent reductions in fitness following introduction (75, 76). Body size is often used as a proxy for fecundity in stickleback (94), which may be particularly important during this phase of establishment and population growth (95). Parasite prevalence could be important as the tapeworm *Schistocephalus solidus* was present in all the source lakes but with great variation in infection rates (52, 96). Populations plagued with higher infection rates during introduction could be at an immediate competitive disadvantage as infection from *S. solidus* decreases growth, fecundity, and fitness (97, 98). Finally, greater genetic diversity within a population might confer greater fitness and establishment success (78, 79). Methods for quantifying each of these characteristics are described in the Supporting Information.

To test if these traits are predictive of population success, we correlated each against the estimated marginal mean effect size of each source population level from the LMMs, estimated through the *emmeans* (v1.11.2-8) package and averaged across the three models of differing environmental distance metrics. Since were interested in how population characteristics might contribute to immediate and longer term success, we regressed population characteristics against effect sizes from both the year one models and the net change models, separately. As regression of population-level effect sizes and traits requires replication at the level of source population, we were limited in power by having only eight replicate source populations, so we consider this analysis to be explorative. As such, we could not include all traits as predictor variables in a single model due to over-fitting; instead, we tested each trait as a predictor individually in separate linear models (Table S8). To account for multiple testing, p-values were adjusted using a Benjamini-Hochberg correction.

## Acknowledgments

Our field sites reside on the traditional and current lands of Dena’ina Peoples, as well as private landowners, and we are grateful for the continued support of all those who allow us to work there. Fieldwork was made possible by collaboration with the Alaska Department of Fish and Game, the Salamatof Native Association, and the Kenai National Wildlife Refuge, as well as the help of many individuals who we list in the Supporting Information. Finally, we thank Antoine Paccard, Ariane Boisclair, and Lena Li Chun Fong of the McGill Genome Centre for collaboration with DNA extraction and genotyping.

