## Supporting Information for "Intrinsic fitness differences outweigh environmental matching in shaping introduction outcomes in nature"

**Supporting Informating**

**Supplementary Tables**

**Table S1:** Summary of source and recipient lakes. For source lakes, ecotype describes the ecotype of the population according to morphology; for recipient lakes, ecotype describes the ecotype of populations introduced to that lake. Environmental and ecological variation among lakes is plotted in Figure S1.

| **Abbr.** | **Name** | **Type** | **Ecotype** | **Region** | **Coordinates** |
| --- | --- | --- | --- | --- | --- |
| FG | Finger Lake | Source | Benthic | Mat-Su | 61.606 N, 149.279 W |
| TL | Tern Lake | Source | Benthic | Kenai | 60.533 N, 149.550 W |
| WB | Walby Lake | Source | Benthic | Mat-Su | 61.620 N, 149.213 W |
| WT | Watson Lake | Source | Benthic | Kenai | 60.539 N, 150.467 W |
| LG | Long Lake | Source | Limnetic | Mat-Su | 61.578 N, 149.764 W |
| SL | Spirit Lake | Source | Limnetic | Kenai | 60.593 N, 150.986 W |
| SR | South Rolly Lake | Source | Limnetic | Mat-Su | 61.667 N, 150.138 W |
| WK | Wik Lake | Source | Limnetic | Kenai | 60.720 N, 151.250 W |
| CC | CC Lake | Recipient | Benthic | Kenai | 60.422 N, 151.195 W |
| LL | Leisure Lake | Recipient | Benthic | Kenai | 60.415 N, 151.210 W |
| LP | Leisure Pond | Recipient | Benthic | Kenai | 60.419 N, 151.207 W |
| CL | Crystal Lake | Recipient | Limnetic | Kenai | 60.424 N, 151.194 W |
| FL | Fred’s Lake | Recipient | Limnetic | Kenai | 60.423 N, 151.200 W |
| HL | Hope Lake | Recipient | Limnetic | Kenai | 60.422 N, 151.188 W |
| RL | Ranchero Lake | Recipient | Limnetic | Kenai | 60.423 N, 151.183 W |
| GL | G Lake | Recipient | Both | Kenai | 60.430 N, 151.177 W |
| LO | Loon Lake | Recipient | Both | Kenai | 60.520 N, 151.051 W |

**Table S2**: Stocking proportions. The number of fish from each source population (columns) introduced to each recipient lake (rows) after accounting for mortalities during translocation.

|  | **LG** | **SL** | **SR** | **WK** | **FG** | **TL** | **WB** | **WT** | **Total** |
| --- | --- | --- | --- | --- | --- | --- | --- | --- | --- |
| **CL** | 366 | 393 | 388 | 270 | 0 | 0 | 0 | 0 | 1417 |
| **FL** | 99 | 100 | 90 | 97 | 0 | 0 | 0 | 0 | 386 |
| **HL** | 430 | 452 | 400 | 287 | 0 | 0 | 0 | 0 | 1569 |
| **RL** | 193 | 199 | 180 | 151 | 0 | 0 | 0 | 0 | 723 |
| **CC** | 0 | 0 | 0 | 0 | 202 | 179 | 198 | 202 | 781 |
| **LL** | 0 | 0 | 0 | 0 | 359 | 243 | 391 | 399 | 1392 |
| **LP** | 0 | 0 | 0 | 0 | 94 | 95 | 100 | 104 | 393 |
| **LO** | 245 | 295 | 242 | 132 | 288 | 130 | 282 | 287 | 1901 |
| **GL** | 0 | 500 | 500 | 495 | 500 | 500 | 500 | 500 | 3495 |

**Table S3:** SNPs unique to each source population that were used for ancestry inference. Positions correspond to the gasAcu1 reference genome (1).

| **Limnetic Assay** | | | | | **Benthic Assay** | | | | |
| --- | --- | --- | --- | --- | --- | --- | --- | --- | --- |
| **Pop.** | **Chr.** | **Pos.** | **Base** | **Freq.** | **Pop.** | **Chr.** | **Pos.** | **Base** | **Freq.** |
| LG | chrII | 15917728 | T | 0.76 | FG | chrI | 15704205 | T | 0.66 |
| LG | chrIII | 823613 | A | 0.76 | FG | chrII | 8998980 | T | 0.78 |
| LG | chrIII | 9359110 | A | 0.68 | FG | chrII | 2892706 | C | 0.76 |
| LG | chrIV | 20067576 | A | 0.85 | FG | chrIII | 11904068 | C | 0.74 |
| LG | chrIV | 13299762 | T | 0.75 | FG | chrIV | 31662263 | T | 0.65 |
| LG | chrV | 1753768 | A | 0.66 | FG | chrIX | 2916678 | T | 0.74 |
| LG | chrVI | 7405819 | A | 0.64 | FG | chrV | 8194471 | C | 0.77 |
| LG | chrVII | 21415801 | T | 0.69 | FG | chrVI | 9113425 | T | 0.61 |
| LG | chrVIII | 14289522 | T | 0.71 | FG | chrVII | 20096576 | A | 0.62 |
| LG | chrX | 6603858 | T | 0.68 | FG | chrVIII | 7909323 | T | 0.49 |
| LG | chrXI | 16193369 | C | 0.91 | FG | chrX | 13397132 | A | 0.58 |
| LG | chrXI | 3081566 | A | 0.80 | FG | chrXII | 2312380 | A | 0.58 |
| LG | chrXII | 2375936 | A | 0.72 | FG | chrXIII | 3776929 | A | 0.53 |
| LG | chrXIII | 8095355 | T | 0.73 | FG | chrXIV | 13767296 | T | 0.66 |
| LG | chrXIV | 2478199 | A | 0.61 | FG | chrXV | 12722008 | A | 0.93 |
| LG | chrXV | 14863116 | T | 0.56 | FG | chrXV | 13373000 | A | 0.80 |
| LG | chrXVII | 9235480 | T | 0.61 | FG | chrXVI | 16659604 | C | 0.70 |
| LG | chrXVIII | 5379465 | T | 0.67 | FG | chrXVII | 9976657 | A | 0.69 |
| LG | chrXX | 3621731 | C | 0.69 | FG | chrXVIII | 6306103 | A | 0.85 |
| LG | chrXXI | 6464598 | A | 0.58 | FG | chrXVIII | 14565840 | A | 0.78 |
| SL | chrI | 6553442 | T | 0.90 | FG | chrXX | 19635221 | A | 0.86 |
| SL | chrII | 7692517 | T | 0.97 | FG | chrXX | 5821398 | A | 0.76 |
| SL | chrII | 7692967 | A | 0.96 | FG | chrXXI | 10304559 | A | 0.49 |
| SL | chrIII | 9231340 | C | 0.93 | TL | chrI | 20114340 | C | 1.00 |
| SL | chrIV | 14839069 | T | 0.95 | TL | chrI | 20163367 | A | 1.00 |
| SL | chrVI | 8431323 | A | 0.95 | TL | chrII | 21053847 | T | 1.00 |
| SL | chrVI | 8425114 | A | 0.94 | TL | chrIV | 5729611 | T | 1.00 |
| SL | chrVII | 1817761 | A | 1.00 | TL | chrII | 20097809 | A | 1.00 |
| SL | chrVIII | 3643167 | A | 0.90 | TL | chrIX | 348981 | C | 1.00 |
| SL | chrX | 7342470 | A | 0.78 | TL | chrV | 2590697 | T | 1.00 |
| SL | chrXVI | 7290192 | T | 0.91 | TL | chrVI | 8817301 | C | 1.00 |
| SL | chrXI | 931319 | T | 0.98 | TL | chrVII | 1723789 | A | 1.00 |
| SL | chrXIII | 591253 | T | 0.90 | TL | chrX | 3733989 | T | 0.98 |
| SL | chrXIV | 11492271 | T | 0.68 | TL | chrXI | 658255 | T | 0.99 |
| SL | chrXV | 504571 | A | 0.80 | TL | chrXII | 136786 | A | 1.00 |
| SL | chrXVI | 10929953 | A | 0.92 | TL | chrXIII | 915897 | A | 1.00 |
| SL | chrXVII | 7968281 | T | 0.88 | TL | chrXV | 10269146 | A | 0.99 |
| SR | chrI | 26934908 | C | 1.00 | TL | chrXVI | 5926417 | A | 1.00 |
| SR | chrI | 26952877 | A | 1.00 | TL | chrXVII | 8487039 | T | 1.00 |
| SR | chrII | 13580477 | T | 0.92 | TL | chrIII | 130727 | A | 1.00 |
| SR | chrIII | 11145910 | A | 0.78 | TL | chrXVIII | 766384 | C | 0.99 |
| SR | chrIV | 32386335 | A | 0.97 | TL | chrXX | 14862997 | A | 1.00 |
| SR | chrIX | 19733496 | A | 0.93 | TL | chrXXI | 11630457 | A | 0.99 |
| SR | chrV | 9747888 | A | 0.94 | WB | chrI | 11658546 | T | 0.68 |
| SR | chrV | 9747863 | C | 0.94 | WB | chrIII | 6828202 | T | 0.67 |
| SR | chrVI | 15534687 | A | 0.79 | WB | chrIX | 11571874 | A | 0.92 |
| SR | chrVII | 2508228 | A | 1.00 | WB | chrIX | 11569307 | T | 0.91 |
| SR | chrVIII | 18169994 | A | 0.92 | WB | chrV | 2233732 | T | 0.62 |
| SR | chrX | 15470216 | A | 0.84 | WB | chrVI | 11736089 | T | 0.69 |
| SR | chrXI | 8508229 | T | 0.96 | WB | chrVII | 7165503 | T | 0.80 |
| SR | chrIX | 10856659 | T | 0.93 | WB | chrVIII | 5789299 | C | 0.63 |
| SR | chrXII | 13556549 | A | 0.95 | WB | chrX | 651167 | T | 0.63 |
| SR | chrXIII | 3295246 | A | 0.78 | WB | chrXI | 2032558 | A | 0.84 |
| SR | chrXIV | 6944009 | T | 0.89 | WB | chrXIII | 77338 | T | 0.64 |
| SR | chrXV | 6208432 | T | 0.72 | WB | chrXIV | 5133486 | A | 0.61 |
| SR | chrXVI | 11846433 | A | 0.84 | WB | chrXV | 10053758 | A | 0.63 |
| SR | chrXVII | 6014009 | T | 0.95 | WB | chrXVI | 11188280 | A | 0.93 |
| SR | chrXVIII | 6861019 | T | 0.77 | WB | chrXVII | 4145810 | T | 0.67 |
| SR | chrXX | 3592900 | A | 0.99 | WB | chrXVIII | 4068044 | T | 0.91 |
| SR | chrXX | 3813198 | T | 0.99 | WB | chrXVIII | 4078056 | T | 0.90 |
| SR | chrXXI | 11294121 | T | 0.85 | WB | chrXXI | 11305884 | T | 0.84 |
| WK | chrI | 9646224 | C | 0.88 | WT | chrI | 6268283 | A | 0.74 |
| WK | chrII | 1721718 | A | 0.71 | WT | chrIII | 8273044 | A | 0.63 |
| WK | chrIV | 29635767 | T | 0.95 | WT | chrIV | 25951865 | T | 0.74 |
| WK | chrIX | 11691955 | A | 0.77 | WT | chrIV | 13840483 | A | 0.72 |
| WK | chrV | 9684570 | A | 0.95 | WT | chrV | 7449895 | T | 0.68 |
| WK | chrV | 2195413 | T | 0.88 | WT | chrVII | 27191714 | T | 0.66 |
| WK | chrVI | 2510463 | T | 0.82 | WT | chrVIII | 3764596 | A | 0.79 |
| WK | chrVI | 10480595 | T | 0.80 | WT | chrX | 3240678 | A | 0.82 |
| WK | chrVII | 7026346 | A | 0.88 | WT | chrX | 3233056 | A | 0.82 |
| WK | chrVIII | 6263140 | T | 0.68 | WT | chrXI | 11603955 | T | 0.73 |
| WK | chrX | 6995177 | A | 0.75 | WT | chrXII | 9432933 | A | 0.71 |
| WK | chrXI | 12958240 | T | 0.65 | WT | chrXIII | 7970424 | A | 0.70 |
| WK | chrXII | 13741528 | T | 0.61 | WT | chrXX | 3577020 | A | 0.75 |
| WK | chrXIII | 14435087 | C | 0.70 |  |  |  |  |  |
| WK | chrXIV | 6924894 | T | 0.81 |  |  |  |  |  |
| WK | chrXV | 1193653 | A | 0.70 |  |  |  |  |  |
| WK | chrXVI | 10258483 | T | 1.00 |  |  |  |  |  |
| WK | chrXVII | 7254362 | T | 0.89 |  |  |  |  |  |
| WK | chrXVII | 7010766 | A | 0.87 |  |  |  |  |  |
| WK | chrXVIII | 14389316 | T | 0.71 |  |  |  |  |  |
| WK | chrXX | 13672198 | T | 0.98 |  |  |  |  |  |
| WK | chrXX | 13069503 | A | 0.93 |  |  |  |  |  |
| WK | chrXXI | 2104869 | C | 0.65 |  |  |  |  |  |

**Table S4**: Observed shifts in source population proportions compared to simulations of drift and sampling error. Statistical significance was determined by the proportion of simulated values with an absolute magnitude greater than the observed change.

| **Recipient Lake** | **Source** | **Obs. Δ prop.** | **Null mean Δ prop.** | **Null CI** | **p-value** |
| --- | --- | --- | --- | --- | --- |
| CC | FG | -0.092 | -0.001 | -0.089 - 0.091 | 0.039 |
| CC | TL | -0.195 | 0.001 | -0.079 - 0.091 | 0.001 |
| CC | WB | 0.082 | 0 | -0.084 - 0.096 | 0.082 |
| CC | WT | 0.205 | 0.001 | -0.089 - 0.092 | 0.001 |
| CL | LG | 0.198 | -0.002 | -0.088 - 0.092 | 0.001 |
| CL | SL | -0.148 | 0.001 | -0.087 - 0.093 | 0.003 |
| CL | SR | 0.09 | -0.001 | -0.094 - 0.096 | 0.056 |
| CL | WK | -0.139 | 0.001 | -0.081 - 0.079 | 0.002 |
| FL | LG | 0.207 | -0.003 | -0.096 - 0.104 | 0.001 |
| FL | SL | -0.13 | 0.004 | -0.089 - 0.101 | 0.011 |
| FL | SR | -0.006 | 0.001 | -0.083 - 0.097 | 0.913 |
| FL | WK | -0.071 | -0.002 | -0.091 - 0.099 | 0.151 |
| HL | LG | 0.22 | 0.003 | -0.084 - 0.096 | 0.001 |
| HL | SL | -0.153 | -0.002 | -0.098 - 0.092 | 0.001 |
| HL | SR | -0.024 | 0 | -0.085 - 0.095 | 0.653 |
| HL | WK | -0.043 | -0.001 | -0.073 - 0.077 | 0.306 |
| LL | FG | -0.088 | 0 | -0.088 - 0.092 | 0.066 |
| LL | TL | -0.159 | -0.001 | -0.075 - 0.075 | 0.001 |
| LL | WB | 0.211 | -0.001 | -0.091 - 0.089 | 0.001 |
| LL | WT | 0.036 | 0.002 | -0.087 - 0.093 | 0.488 |
| LO | FG | -0.123 | 0.001 | -0.061 - 0.079 | 0.003 |
| LO | TL | -0.05 | 0 | -0.038 - 0.052 | 0.051 |
| LO | WB | -0.11 | 0 | -0.068 - 0.082 | 0.003 |
| LO | WT | 0.073 | -0.001 | -0.071 - 0.069 | 0.037 |
| LO | LG | 0.137 | -0.001 | -0.059 - 0.071 | 0.001 |
| LO | SL | -0.047 | 0.001 | -0.075 - 0.085 | 0.191 |
| LO | SR | 0.19 | 0.001 | -0.057 - 0.073 | 0.001 |
| LO | WK | -0.069 | 0 | -0.049 - 0.051 | 0.009 |
| LP | FG | -0.114 | -0.003 | -0.089 - 0.091 | 0.01 |
| LP | TL | -0.195 | -0.001 | -0.092 - 0.099 | 0.001 |
| LP | WB | 0.048 | 0.001 | -0.094 - 0.106 | 0.318 |
| LP | WT | 0.261 | 0.003 | -0.085 - 0.105 | 0.001 |
| RL | LG | 0.195 | 0.002 | -0.087 - 0.103 | 0.001 |
| RL | SL | -0.103 | -0.001 | -0.095 - 0.095 | 0.04 |
| RL | SR | -0.004 | -0.001 | -0.079 - 0.091 | 0.916 |
| RL | WK | -0.088 | 0 | -0.079 - 0.081 | 0.046 |
| GL | FG | -0.038 | 0 | -0.063 - 0.067 | 0.25 |
| GL | TL | -0.123 | -0.002 | -0.063 - 0.067 | 0.001 |
| GL | WB | 0.008 | -0.001 | -0.063 - 0.077 | 0.763 |
| GL | WT | 0.028 | 0.001 | -0.063 - 0.077 | 0.4 |
| GL | SL | -0.015 | 0 | -0.063 - 0.077 | 0.671 |
| GL | SR | 0.124 | 0.001 | -0.063 - 0.077 | 0.001 |
| GL | WK | 0.016 | 0.001 | -0.072 - 0.068 | 0.664 |

**Table S5**: Models of source success. Results from linear mixed models of source success over three timespans. Five alternative measures of environmental similarity were used. All models included source population identity and initial frequency as fixed effects and recipient lake as a random effect.

| **Time** | **Env. Dist.** | **Parameter** | **Effect size (β)** | **SE** | **CI** | **t-value** | **df** | **p-value** | **p_adj_** |
| --- | --- | --- | --- | --- | --- | --- | --- | --- | --- |
| 1 | RLA+calc | Env. Dist. | -0.106 | 0.109 | -0.422 - 0.21 | 26 | -0.692 | 0.495 | 0.675 |
| 1 | RLA+calc | Init. Prop. | -0.076 | 0.157 | -0.527 - 0.374 | 26 | -0.350 | 0.729 | 0.810 |
| 1 | prey | Env. Dist. | -0.050 | 0.080 | -0.295 - 0.195 | 26 | -0.420 | 0.678 | 0.810 |
| 1 | prey | Init. Prop. | -0.061 | 0.157 | -0.511 - 0.389 | 26 | -0.279 | 0.782 | 0.814 |
| 1 | all | Env. Dist. | -0.116 | 0.118 | -0.489 - 0.257 | 26 | -0.641 | 0.527 | 0.688 |
| 1 | all | Init. Prop. | -0.059 | 0.156 | -0.507 - 0.388 | 26 | -0.274 | 0.787 | 0.814 |
| 2 | RLA+calc | Env. Dist. | 0.222 | 0.091 | -0.159 - 0.602 | 26 | 1.202 | 0.240 | 0.424 |
| 2 | RLA+calc | Init. Prop. | -0.404 | 0.133 | -0.956 - 0.148 | 26 | -1.509 | 0.143 | 0.347 |
| 2 | prey | Env. Dist. | 0.139 | 0.068 | -0.161 - 0.44 | 26 | 0.958 | 0.347 | 0.496 |
| 2 | prey | Init. Prop. | -0.381 | 0.135 | -0.942 - 0.18 | 26 | -1.401 | 0.173 | 0.347 |
| 2 | all | Env. Dist. | 0.176 | 0.101 | -0.285 - 0.637 | 26 | 0.786 | 0.439 | 0.549 |
| 2 | all | Init. Prop. | -0.390 | 0.136 | -0.953 - 0.173 | 26 | -1.429 | 0.165 | 0.347 |
| Net | RLA+calc | Env. Dist. | -0.011 | 0.100 | -0.29 - 0.268 | 33 | -0.079 | 0.937 | 0.989 |
| Net | RLA+calc | Init. Prop. | 0.259 | 0.137 | -0.084 - 0.602 | 33 | 1.542 | 0.133 | 0.221 |
| Net | prey | Env. Dist. | -0.019 | 0.073 | -0.222 - 0.184 | 33 | -0.188 | 0.852 | 0.946 |
| Net | prey | Init. Prop. | 0.257 | 0.133 | -0.076 - 0.59 | 33 | 1.574 | 0.125 | 0.221 |
| Net | all | Env. Dist. | -0.082 | 0.103 | -0.369 - 0.204 | 33 | -0.585 | 0.562 | 0.734 |
| Net | all | Init. Prop. | 0.249 | 0.131 | -0.079 - 0.578 | 33 | 1.549 | 0.131 | 0.221 |

**Table S6**: Effects of source identity on population success across all models, estimated through Type III F-tests.

| **Time** | **Env. Dist.** | **SS** | **MS** | **df_1_** | **df_2_** | **F-value** | **p-value** | **p_adj_** |
| --- | --- | --- | --- | --- | --- | --- | --- | --- |
| 1 | RLA+calc | 4.171 | 0.596 | 7 | 26 | 3.425 | 0.010 | 0.015 |
| 1 | prey | 5.348 | 0.764 | 7 | 26 | 4.341 | 0.003 | 0.008 |
| 1 | all | 2.804 | 0.401 | 7 | 26 | 2.296 | 0.058 | 0.058 |
| 2 | RLA+calc | 3.416 | 0.488 | 7 | 26 | 3.987 | 0.004 | 0.006 |
| 2 | prey | 3.339 | 0.477 | 7 | 26 | 3.823 | 0.006 | 0.006 |
| 2 | all | 3.461 | 0.494 | 7 | 26 | 3.918 | 0.005 | 0.006 |
| net | RLA+calc | 8.221 | 1.174 | 7 | 33 | 6.704 | 0.000 | 0.000 |
| net | prey | 8.736 | 1.248 | 7 | 33 | 7.130 | 0.000 | 0.000 |
| net | all | 4.761 | 0.680 | 7 | 33 | 3.922 | 0.003 | 0.003 |

**Table S7**: Estimated marginal means (EMMs) effect sizes of each source population in each source population success model.

| **Time** | **Env. Dist.** | **Source** | **EMM** | **SE** | **df** | **CI** |
| --- | --- | --- | --- | --- | --- | --- |
| 1 | RLA+calc | FG | -0.537 | 0.241 | 24.842 | -1.033 - -0.04 |
| 1 | prey | FG | -0.589 | 0.222 | 25.128 | -1.047 - -0.131 |
| 1 | all | FG | -0.542 | 0.242 | 24.192 | -1.04 - -0.043 |
| 1 | RLA+calc | LG | 0.378 | 0.208 | 25.956 | -0.05 - 0.806 |
| 1 | prey | LG | 0.408 | 0.201 | 25.960 | -0.005 - 0.822 |
| 1 | all | LG | 0.340 | 0.235 | 23.849 | -0.145 - 0.824 |
| 1 | RLA+calc | SL | 0.046 | 0.229 | 25.813 | -0.424 - 0.516 |
| 1 | prey | SL | 0.068 | 0.225 | 25.563 | -0.394 - 0.53 |
| 1 | all | SL | 0.086 | 0.223 | 25.479 | -0.373 - 0.545 |
| 1 | RLA+calc | SR | -0.117 | 0.192 | 25.915 | -0.513 - 0.278 |
| 1 | prey | SR | -0.094 | 0.191 | 25.954 | -0.488 - 0.299 |
| 1 | all | SR | -0.150 | 0.208 | 24.711 | -0.579 - 0.279 |
| 1 | RLA+calc | TL | -1.423 | 0.304 | 25.007 | -2.05 - -0.796 |
| 1 | prey | TL | -1.466 | 0.297 | 24.610 | -2.078 - -0.854 |
| 1 | all | TL | -1.369 | 0.342 | 25.740 | -2.072 - -0.665 |
| 1 | RLA+calc | WB | 0.156 | 0.240 | 25.775 | -0.336 - 0.649 |
| 1 | prey | WB | 0.110 | 0.227 | 25.953 | -0.357 - 0.578 |
| 1 | all | WB | 0.106 | 0.225 | 25.998 | -0.356 - 0.569 |
| 1 | RLA+calc | WK | -0.479 | 0.276 | 25.870 | -1.046 - 0.089 |
| 1 | prey | WK | -0.436 | 0.264 | 25.255 | -0.981 - 0.108 |
| 1 | all | WK | -0.405 | 0.263 | 25.160 | -0.946 - 0.137 |
| 1 | RLA+calc | WT | 0.235 | 0.247 | 25.804 | -0.272 - 0.742 |
| 1 | prey | WT | 0.228 | 0.249 | 25.810 | -0.283 - 0.74 |
| 1 | all | WT | 0.182 | 0.264 | 25.754 | -0.362 - 0.725 |
| 2 | RLA+calc | FG | -0.228 | 0.203 | 24.667 | -0.647 - 0.191 |
| 2 | prey | FG | -0.162 | 0.189 | 24.899 | -0.552 - 0.227 |
| 2 | all | FG | -0.194 | 0.207 | 24.029 | -0.621 - 0.233 |
| 2 | RLA+calc | LG | 0.448 | 0.218 | 25.921 | -0.001 - 0.897 |
| 2 | prey | LG | 0.388 | 0.214 | 25.999 | -0.052 - 0.829 |
| 2 | all | LG | 0.459 | 0.234 | 25.264 | -0.022 - 0.941 |
| 2 | RLA+calc | SL | -0.481 | 0.174 | 25.175 | -0.839 - -0.123 |
| 2 | prey | SL | -0.523 | 0.167 | 25.756 | -0.867 - -0.18 |
| 2 | all | SL | -0.552 | 0.168 | 25.782 | -0.897 - -0.207 |
| 2 | RLA+calc | SR | 0.301 | 0.160 | 25.969 | -0.029 - 0.63 |
| 2 | prey | SR | 0.267 | 0.160 | 25.979 | -0.063 - 0.597 |
| 2 | all | SR | 0.330 | 0.177 | 24.842 | -0.035 - 0.694 |
| 2 | RLA+calc | TL | -0.404 | 0.274 | 25.912 | -0.968 - 0.159 |
| 2 | prey | TL | -0.311 | 0.264 | 26.000 | -0.853 - 0.232 |
| 2 | all | TL | -0.407 | 0.296 | 24.977 | -1.017 - 0.203 |
| 2 | RLA+calc | WB | -0.088 | 0.199 | 25.938 | -0.497 - 0.322 |
| 2 | prey | WB | -0.039 | 0.193 | 25.862 | -0.436 - 0.358 |
| 2 | all | WB | -0.025 | 0.192 | 25.564 | -0.421 - 0.371 |
| 2 | RLA+calc | WK | -0.518 | 0.204 | 25.365 | -0.938 - -0.098 |
| 2 | prey | WK | -0.545 | 0.201 | 24.884 | -0.959 - -0.131 |
| 2 | all | WK | -0.591 | 0.200 | 24.536 | -1.002 - -0.179 |
| 2 | RLA+calc | WT | 0.374 | 0.199 | 25.121 | -0.036 - 0.783 |
| 2 | prey | WT | 0.369 | 0.200 | 25.069 | -0.043 - 0.781 |
| 2 | all | WT | 0.409 | 0.214 | 25.997 | -0.03 - 0.848 |
| Net | RLA+calc | FG | -0.634 | 0.219 | 31.436 | -1.081 - -0.187 |
| Net | prey | FG | -0.632 | 0.199 | 31.868 | -1.037 - -0.227 |
| Net | all | FG | -0.579 | 0.220 | 30.697 | -1.029 - -0.129 |
| Net | RLA+calc | LG | 0.480 | 0.203 | 32.781 | 0.068 - 0.893 |
| Net | prey | LG | 0.484 | 0.198 | 32.996 | 0.08 - 0.888 |
| Net | all | LG | 0.428 | 0.221 | 30.821 | -0.023 - 0.88 |
| Net | RLA+calc | SL | -0.638 | 0.199 | 33.000 | -1.043 - -0.232 |
| Net | prey | SL | -0.637 | 0.195 | 32.914 | -1.035 - -0.24 |
| Net | all | SL | -0.634 | 0.194 | 32.825 | -1.029 - -0.239 |
| Net | RLA+calc | SR | 0.261 | 0.179 | 32.636 | -0.103 - 0.625 |
| Net | prey | SR | 0.263 | 0.173 | 32.984 | -0.088 - 0.615 |
| Net | all | SR | 0.220 | 0.190 | 31.736 | -0.167 - 0.606 |
| Net | RLA+calc | TL | -1.132 | 0.278 | 32.323 | -1.697 - -0.566 |
| Net | prey | TL | -1.135 | 0.273 | 31.968 | -1.692 - -0.579 |
| Net | all | TL | -1.067 | 0.298 | 32.953 | -1.674 - -0.46 |
| Net | RLA+calc | WB | -0.106 | 0.223 | 32.229 | -0.561 - 0.349 |
| Net | prey | WB | -0.106 | 0.203 | 32.795 | -0.519 - 0.307 |
| Net | all | WB | -0.098 | 0.199 | 32.981 | -0.503 - 0.308 |
| Net | RLA+calc | WK | -0.446 | 0.242 | 32.501 | -0.939 - 0.046 |
| Net | prey | WK | -0.449 | 0.218 | 32.873 | -0.892 - -0.006 |
| Net | all | WK | -0.446 | 0.208 | 32.176 | -0.871 - -0.022 |
| Net | RLA+calc | WT | 0.220 | 0.212 | 32.623 | -0.212 - 0.652 |
| Net | prey | WT | 0.219 | 0.211 | 32.600 | -0.21 - 0.648 |
| Net | all | WT | 0.190 | 0.216 | 32.642 | -0.25 - 0.63 |

**Table S8**: Linear models of population characteristics on the mean estimated marginal mean effect size of each source in the source population success models.

| **Time** | **Predictor** | **Effect size (β)** | **R^2^** | **Adj. R^2^** | **p-value** | **p_adj_** |
| --- | --- | --- | --- | --- | --- | --- |
| 1 | Genetic diversity | 0.487 | 0.237 | 0.110 | 0.221 | 0.442 |
| 1 | Mortality rate | -0.665 | 0.442 | 0.349 | 0.072 | 0.288 |
| 1 | Median body size | 0.141 | 0.020 | -0.144 | 0.739 | 0.739 |
| 1 | Parasite prevalence | -0.230 | 0.053 | -0.105 | 0.583 | 0.739 |
| Net | Genetic diversity | 0.431 | 0.185 | 0.050 | 0.287 | 0.586 |
| Net | Mortality rate | -0.285 | 0.081 | -0.072 | 0.493 | 0.586 |
| Net | Median body size | 0.229 | 0.052 | -0.106 | 0.586 | 0.586 |
| Net | Parasite prevalence | -0.285 | 0.081 | -0.072 | 0.493 | 0.586 |

**Table S9:** Estimates of genetic diversity for each source population. Only nucleotide diversity was used in the models reported in Table S8.

| Source population | Ecotype | Nucleotide diversity (π) | Watterson’s estimator (θ_w_) | Tajima’s D |
| --- | --- | --- | --- | --- |
| FG | Benthic | 0.0012 | 0.0033 | -5.267 |
| TL | Benthic | 0.0015 | 0.0048 | -5.627 |
| WB | Benthic | 0.0029 | 0.0062 | -4.366 |
| WT | Benthic | 0.0016 | 0.0041 | -4.911 |
| LG | Limnetic | 0.0023 | 0.0051 | -4.539 |
| SL | Limnetic | 0.0016 | 0.0040 | -4.963 |
| SR | Limnetic | 0.0016 | 0.0044 | -5.235 |
| WK | Limnetic | 0.0018 | 0.0049 | -5.095 |

**Supplementary Figures**

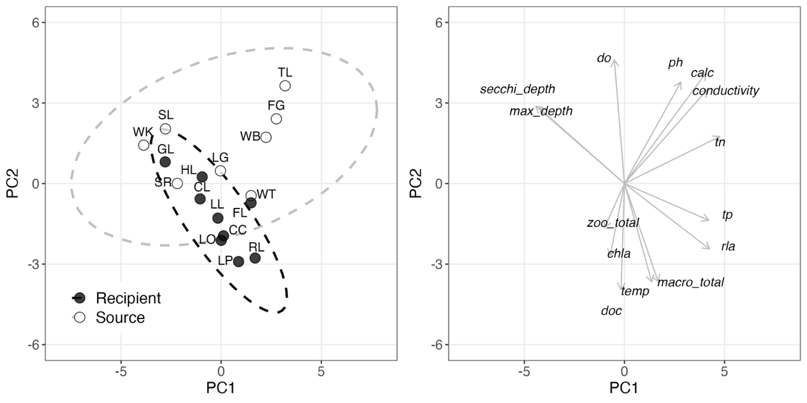

**Figure S1**: PCA of all environmental and ecological measures. Left shows the source and recipient lakes (abbreviations and details in Table S1). Right shows loadings of the following variables: dissolved oxygen (do), pH, calcium concentration (calc), specific conductivity, total nitrogen (tn), total phosporous (tp), relative littoral area (rla), total abundance of key macroinvertebrate taxa (macro_total), temperature (temp), dissolved organic carbon (doc), chlorophyll *a* (chla), total abundance of key zooplankton taxa (zoo_total), maximum depth, and Secchi depth. Methods for measuring each variable are in Supplementary Methods.

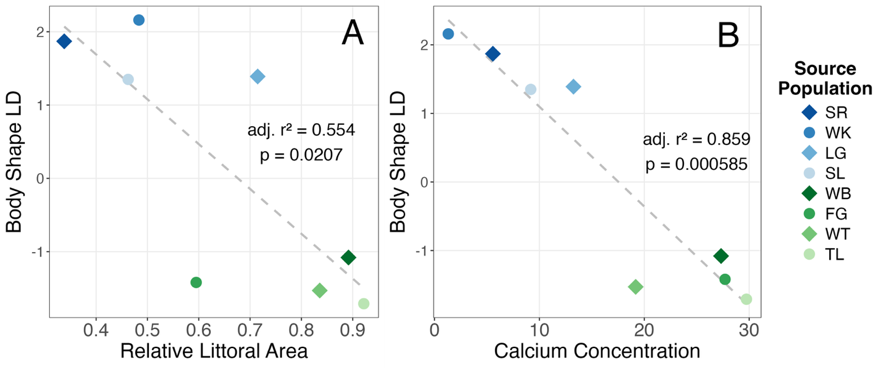

**Figure S2**: Environmental correlations: **(A)** Relative littoral area and **(B)** calcium concentration regressed against a population-mean linear discriminant (LD) score of body shape (from geometric morphometrics) of each source population, from Hendry et al. (2024).

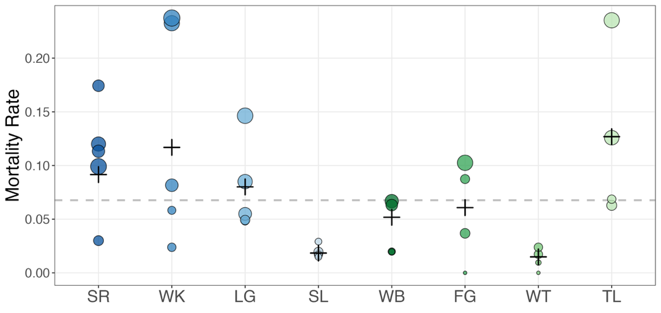

**Figure S3**: Mortality during translocation: Points represent mortality rates for each source by recipient lake pair during the translocation process, scaled by size according to the absolute number of mortalities. Crossbars represent the combined morality rate for each source population, and the dashed line represents the overall mortality rate.

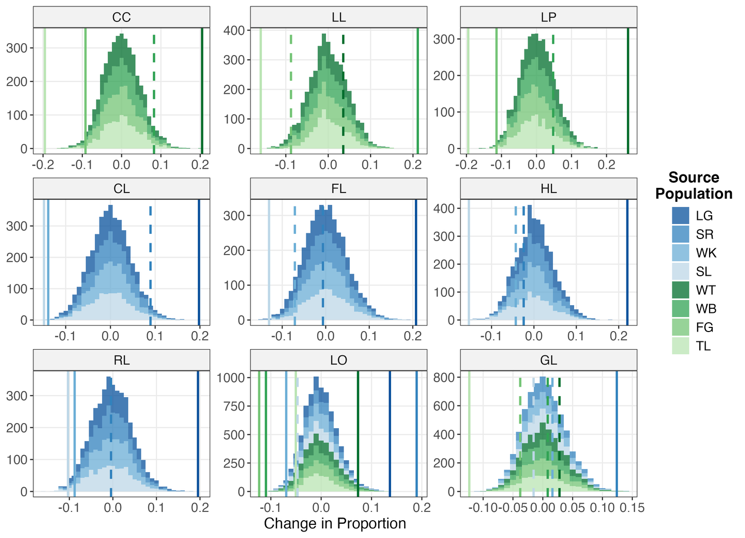

**Figure S4**: Neutral expectations of changes in source population contribution according to parameterized simulations of exponential growth. Histograms show the distribution of expected changes, while the vertical lines represent the observed changes; solid lines are outside the 95% confidence interval for that source population in that lake, dotted lines are within the interval.

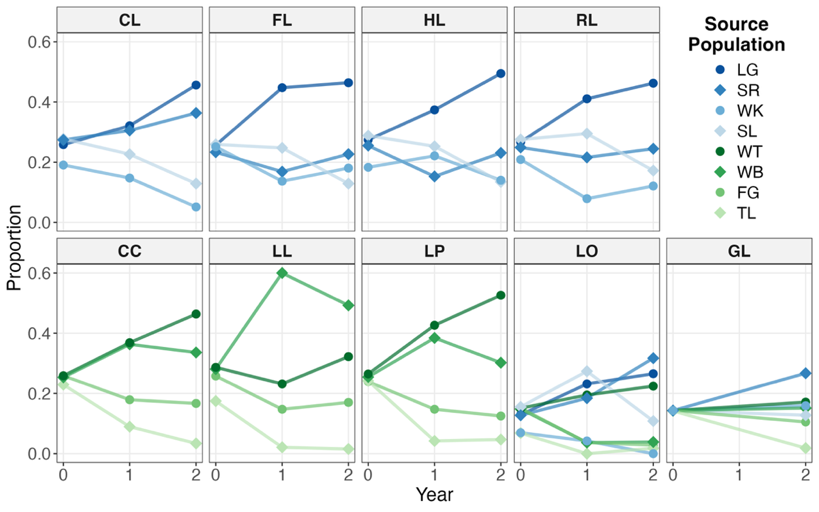

**Figure S5**: Proportional contribution of each source population in each lake over time, showing the same trends as Figure 2, but with raw proportions rather than changes with respect to initial stocking proportions.

**Supplementary Methods**

Environmental and ecological measures

*Environmental measures*

Environmental variables were measured according to the following protocols, as described in (2). We measured dissolved oxygen (DO; mg/L), pH, water temperature (°C), and conductivity (μS/cm2) using a Professional Plus Model YSI multi-parameter sonde (model 10,102,030; Yellow Springs Inc.) at the deepest point of each lake in June 2018 and June 2019 – the values we used were those at 0.5m depth at that point. Secchi depth (m) was estimated by lowering a Secchi disk on the shaded side of boat. We collected water samples to quantify dissolved organic carbon (DOC; mg/L), total nitrogen (TN; mg/L), total phosphorous (TP; μg/L), chlorophyll a (chl-a; μg/L), and dissolved calcium (Ca; mg/L). DOC, TN, TP, and chl-a samples were analyzed at the GRIL-Université du Québec à Montréal (UQAM) analytical laboratory. DOC concentrations of 0.45 μm filtered samples (surfactant-free membrane filters) were measured after acidification (phosphoric acid 5%) followed by sodium persulfate oxidation using a 1010 TOC analyzer (O.I. Analytical, College Station, TX, USA). The absorption at 440 nm (CDOM), used as a measure of watercolor, was measured on water samples with a 2 cm quartz cuve in a BiochromUltrospec® 2100 pro spectrofluorometer (3). We quantified total nitrogen (TN) with a continuous flow analyzer (OI Analytical Flow Solution 3100 ©) using an alkaline persulfate digestion method, coupled with a cadmium reactor, following a standard protocol (4). We quantified total phosphorus (TP) spectrophotometrically on the same machine by the molybdenum blue method after persulfate digestion (5). Chl-a was quantified by passing samples through glass fiber filters (Whatman GF/F), extraction of the Chl-a in hot ethanol, and measuring the chlorophyll spectrophotometrically on a “BiochromUltrospec” 2100 pro with a 10-cm quartz cuvette (6, 7). Water calcium concentrations were analyzed with a Thermo ICAP-6300 Inductively Coupled Argon Plasma—Optical Emission Spectrometer (ICP-OES) following protocols described by US EPA (1997) at the University of Alberta Biogeochemical Analytical Service Laboratory (U of A—BASL; Edmonton, Alberta, Canada).

*Ecological measures*

Crustacean zooplankton were collected by whole water column vertical tows with a 35-cm diameter Wisconsin net from 0.5 m off the bottom to the surface. The zooplankton were sampled from four sampling stations along an open water transect across each lake. The crustacean zooplankton were anaesthetized with bromoseltzer, preserved with 95% ethanol, and they were brought back to the laboratory at the Université du Québec à Montréal (UQAM) for identification. These four samples were subsequently pooled to a single sample per lake for identification and enumeration in the laboratory. The crustacean zooplankton were identified to species level with a high- resolution dissecting microscope (6.3-126x; SZ2-IL-ST; Olympus SZ) and species were enumerated for the number of individuals per L. Crustacean zooplankton were counted using a protocol that targeted mature individuals that could be identified unambiguously to species, as well as to detect rare species (9). Subsamples (10 mL) were taken from a standardised 50 mL sample volume, until at least 200 individuals of the most abundant species were counted (excluding copepodids and nauplii), or a total of 1000 individuals were counted (excluding copepodids and nauplii), or the total sample was counted. Taxonomic keys included (10), (11), (12), (13), (14), (15).

Nearshore littoral macroinvertebrate communities were collected semi-quantitatively with a D-frame kick net at four (2018) to eight (2019) sampling stations located haphazardly around the lake. We increased the number of sampling stations to eight stations per lake in 2019 to improve the capture of spatial variation in macroinvertebrates within each lake (approx. 2 m^2 area sampled per station). Nearshore littoral macroinvertebrate communities were collected with a D-frame kick net by the sweep method (“Kick & Sweep”) with a 500 μm “D-net” as recommended by the Ontario Benthos Biomonitoring Network (16) on a surface of approximatively 2 m^2^. Samples were concentrated with a 500-μm sieve, preserved in 95% ethanol, and they were brought back to the laboratory at the Université du Québec à Montréal (UQAM) for identification. Identification was done up to the Family level following taxonomic keys (17, 18). All macroinvertebrate samples were identified using SZX10 stereo microscopes (Olympus) with varying magnification (x6.3 - x10). For each sample, 100mL sub-sample was taken and counted until the 100th individual was reached. If the 100th individual was not reached within the initial 100 mL, an additional 100 mL was counted. When the 100^th^ individual was reached, the remaining part of the sub-sample was counted, and the total sub-sampled volume calculated. A ratio was then calculated between the total sub-sampled volume and the sample total volume to estimate the taxon-specific abundance of macroinvertebrates. The density of macroinvertebrates per m^2^ was averaged for the 8 samples collected per lake to obtain total macroinvertebrate density per lake.

Measuring population characteristics

*Median body size*

While fish were being sampled for transplant in 2019, an additional 100 fish from each source population were sampled and euthanized. Using this additional sample of fish, the median standard length (measured by digital calipers to 0.01mm) was calculated for each source population.

*Parasite prevalence*

Fish from the additional sample at the time of transplant were dissected and the number and mass of *Schistocephalus solidus* present in each fish was recorded. For our models, we use infection prevalence (the proportion of fish from each population with at least one *S. solidus* infection).

*Genetic diversity*

As described in Weber et al. (2022), source populations were sequenced as pools comprised of 100 fish each that were sampled in 2018. We accessed the raw reads from this sequencing and trimmed them using *fastp* (v.0.24.0), removing reads with Phred scores less than 20 or lengths shorter than 50bp, as well as adapter sequences and polyG tails. We performed quality control on both the raw and trimmed reads using *fastQC* (v.0.12.1) and *MultiQC* (v.1.31). The trimmed reads were then aligned to the University of Georgia *G. aculeatus* reference genome v.5 (20) using *BWA* (v.0.7.18), filtering out reads with a mapping score less than 20. The resulting bam files were sorted and indexed with *Samtools* (v.1.20), and PCR duplicates were marked and removed using *Picard* (v.3.1.0). We identified and masked indel regions (5bp above and below each indel) using *PoPoolation2* (21), before converting to a sync file and filtering out positions with a base quality less than 20. Using *grenedalf* v.0.6.3 (22), we estimated three metrics of genetic diversity for each population: nucleotide diversity (π), the proportion of segregating sites (Watterson’s estimator; θ_w_), and Tajima’s D. For each metric, we estimated diversity across the autosomes (removing the sex chromosomes, mitochondrial sequences, and other scaffolds), filtering out positions with a minimum read count less than 2, a minimum depth less than 4, or a maximum depth greater than 200 and averaged estimates across all remaining positions in the genome. In our models of source population characteristics, we only used nucleotide diversity, but all metrics are shown in Table S9.

Extended acknowledgements

We thank the fieldwork crews that conducted the transplant and collected fish in 2019, 2020, 2021, 2022, and 2024, including Ismail Ameen, Luis Baertschi, Tina Barbasch, Steven Bezdecny, Chelsea Bishop, Chad Brock, Sheila Christen, Mariane Daneau-Lamoureux, Victor Frankel, Gregor Fussmann, Alex Gouvin Moffat, Stéphanie Guernon, Rachel Kramp, Kelly Ireland, Matt Josephson, Zuyao Liu, Kathryn Milligan-McClellan, Louis Moisan, Anya Mueller, Kevin Neumann, Michelle Packer, Arshad Padhiar, Sarah Pasqualetti, Allegra Pearce, Christopher Peterson, Hillary Poore, Maria Rodgers, Andrea Roth, Rogini Runghen, Sarah Sanderson, Trey Sasser, Hiranya Sudasinghe, Richard Benjamin Sulser, Saraswathy Vaidyanathan, Matthew Walsh, and Anika Wohlleben.
